# Fronto-Temporal Dysconnectivity and Cortical Excitability in High Schizotypy: Associations with Symptom Dimensions

**DOI:** 10.64898/2026.04.16.718911

**Authors:** Daniel J. Hauke, Galya C. Iseli, Julia Rodriguez Sanchez, James M. Stone, David Coynel, Rick A. Adams, André Schmidt

**Affiliations:** Hawkes Institute, Department of Computer Science, University College London, London, UK; University of Basel, Department of Clinical Research (DKF), University Psychiatric Clinics (UPK), Translational Neurosciences, Basel, Switzerland; Division of Cognitive Neuroscience, Department of Biomedicine, University of Basel, Basel, Switzerland; Research Cluster Molecular and Cognitive Neurosciences, Department of Biomedicine, University of Basel, Basel, Switzerland; School of Health and Medical Sciences, City St George’s, University of London,, London, UK; Brighton and Sussex Medical School, University of Sussex, Brighton, UK; Institute of Cognitive Neuroscience, University College London, London, UK

**Keywords:** schizotypy, E/I balance, effective connectivity, psychosis spectrum, schizophrenia

## Abstract

**Background:** Psychosis has been conceptualised as a continuum extending from healthy individuals with psychotic-like experiences to clinical populations with schizophrenia. It is unclear which biological mechanisms found in chronic schizophrenia extend across the psychosis continuum to healthy individuals with high positive schizotypy (HS). In this study, we used computational modeling to test whether changes in effective connectivity and excitation/inhibition (E/I) balance reported in schizophrenia are also found in HS.

**Methods:** A total of 2425 individuals from the general population were screened for HS. A subset (N=141) was invited for in-depth phenotyping. Resting-state functional magnetic resonance imaging (rsfMRI) and proton magnetic resonance spectroscopy (^1^H-MRS) were recorded in n=69 HS individuals and n=72 group-matched controls with low schizotypy (LS). We used dynamic causal modeling to estimate effective connectivity between bilateral primary auditory cortex (A1), superior temporal gyrus (STG), and inferior frontal gyrus (IFG).

**Results:** Bilateral backward connectivity from IFG to STG was significantly reduced in HS compared to LS. Widespread cortical disinhibition in the auditory cortex-IFG network correlated with more severe positive schizotypy scores and impulsive nonconformity. Reduced excitability in the same network was correlated with stronger cognitive disorganisation.

**Conclusions:** Our results favour a psychosis-continuum hypothesis, suggesting that reduced top-down drive from frontal cortex and compensatory allostatic upregulation of cortical excitability, as observed in chronic schizophrenia, also extend to groups with sub-clinical psychotic symptoms. Frontal cortex dysfunction may serve as a biologically interpretable biomarker of psychosis risk and a target for preventative interventions.

## Introduction

Schizotypy is a multidimensional construct that captures subclinical traits and experiences resembling the core symptom domains of schizophrenia, including unusual perceptual experiences, social withdrawal, cognitive disorganization and impulsive nonconformity (1,2). Rather than representing a discrete diagnostic category, schizotypy has been conceptualized as a continuum of psychosis-proneness, observable across clinical and non-clinical populations (3–6). This dimensional view aligns with the Research Domain Criteria (RDoC) initiative, which advocates for a neuroscience-grounded classification system based on observable behaviour and neurobiological mechanisms rather than traditional diagnostic boundaries (7–9). However, it remains unclear, which biological mechanisms found in schizophrenia cut across the psychosis-spectrum to healthy individuals with high schizotypy and could support such a neuroscience-grounded classification system.

A key biological mechanism that has been put forth as an aetiological explanation for schizophrenia is disruption of the excitation/inhibition (E/I) balance (10,11). Evidence supporting E/I imbalance in schizophrenia comes from genetic (12,13), post-mortem (14,15), electroencephalography (EEG) (16–31), and magnetic resonance spectroscopy (MRS) studies (32–36).

For instance, hippocampal glutamate is increased in individuals at clinical high risk for psychosis (CHR-P) who converted to a psychotic disorder (37,38) and those with poor functional outcomes (38). However, evidence in CHR-P is still inconsistent (32,34,35) with an earlier meta-analysis finding elevated glutamate in medial frontal and not medial temporal regions in CHR-P (34). In chronic patients with schizophrenia, Merritt and colleagues (34) reported elevated glutamate in the medial temporal lobe, which was also found in first-episode psychosis patients by others (36). A more recent mega-analysis additionally found increased glutamate in medial temporal lobe to be correlated with more severe positive symptoms and worse functioning (35).

In terms of electrophysiology, Hauke et al. (20) found that E/I imbalance caused by reduced pyramidal cell excitability could explain reduced event-related potentials across three different auditory paradigms - including the paired-click (or ‘sensory-gating’), mismatch negativity (MMN) and P300 - in a simulation study. In line with this, Adams et al. (16) found that pyramidal cell excitability was consistently reduced (especially in frontal regions) across four different experimental paradigms (including three EEG paradigms and resting-state functional magnetic resonance imaging, rs-fMRI) in patients with chronic schizophrenia. Rodriguez-Sanchez et al. (30) found that pyramidal cell excitability was already reduced in CHR-P individuals who converted to a psychotic disorder within two years, compared to those who remitted, indicating that reduced pyramidal excitability may be a primary pathology of schizophrenia that is already present in the prodromal phase. Interestingly, however, *disinhibition* was associated with more severe positive symptoms in Adams et al. (16) and Rodriguez-Sanchez et al. (30), suggesting that symptoms may result from a compensatory allostatic upregulation of excitability. Importantly, it is yet unknown whether these biological mechanisms extend to healthy individuals with schizotypal traits.

In this study, we use computational modeling of rs-fMRI to compare individuals with high compared to low positive schizotypal traits to investigate this question. Based on Adams et al. (16), we predicted reduced excitability in frontal cortex in individuals with pronounced schizotypal traits, but disinhibition to be associated with more severe psychotic-like experiences.

## Methods

This study analyses data from Iseli et al. (39). Please, see the original study for further details on the protocol and previous findings of the MRS and clinical data. For convenience, here we summarise the most pertinent details for this study below.

### Participants

A total of 2425 healthy individuals from the general population were screened for high positive schizotypy traits measured with the *Unusual Experiences* scale of the German translation of the short Oxford-Liverpool Inventory of Feelings and Experiences (sO-LIFE) (1,2,40). Positive schizotypy as opposed to other subscales were selected to align this study with diagnostic definitions of schizophrenia (41) and the clinical high risk state (42,43), which emphasise this symptom dimension. Based on this screening, *n*=141 participants including *n*=69 with high positive schizotypy (HS; *Unusual Experiences* ≥ 6, i.e. >1.5 SD above the median) and *n*=72 control individuals with low positive schizotypy (LS; *Unusual Experiences* ≤ 1) were invited for in-depth phenotyping and neuroimaging assessments (MRS, rsfMRI). LS were prospectively group-matched to HS in terms of sex, age, smoking status and cannabis consumption (Table 1). Individuals with an acute psychiatric disorder diagnosis or first-degree family history of schizophrenia, schizoaffective disorders, bipolar disorder, autism spectrum disorder or suicide were excluded from the study. This study was approved by the Ethics Committee of Northwestern and Central Switzerland (EKNZ Basel, BASEC number 2021-00933) and was conducted in accordance with the Declaration of Helsinki. All participants provided written informed consent prior to study enrolment.

**Table 1.**
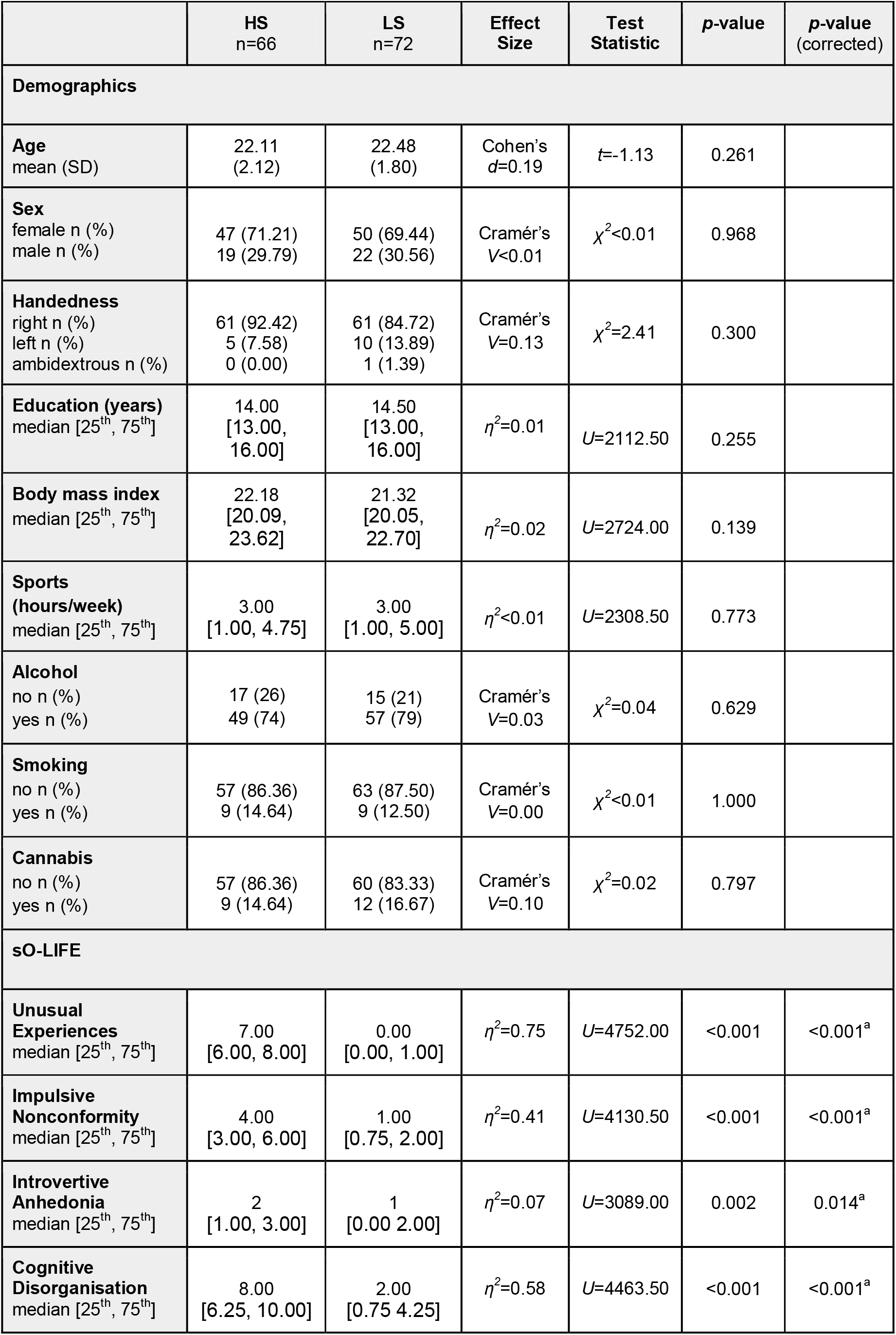
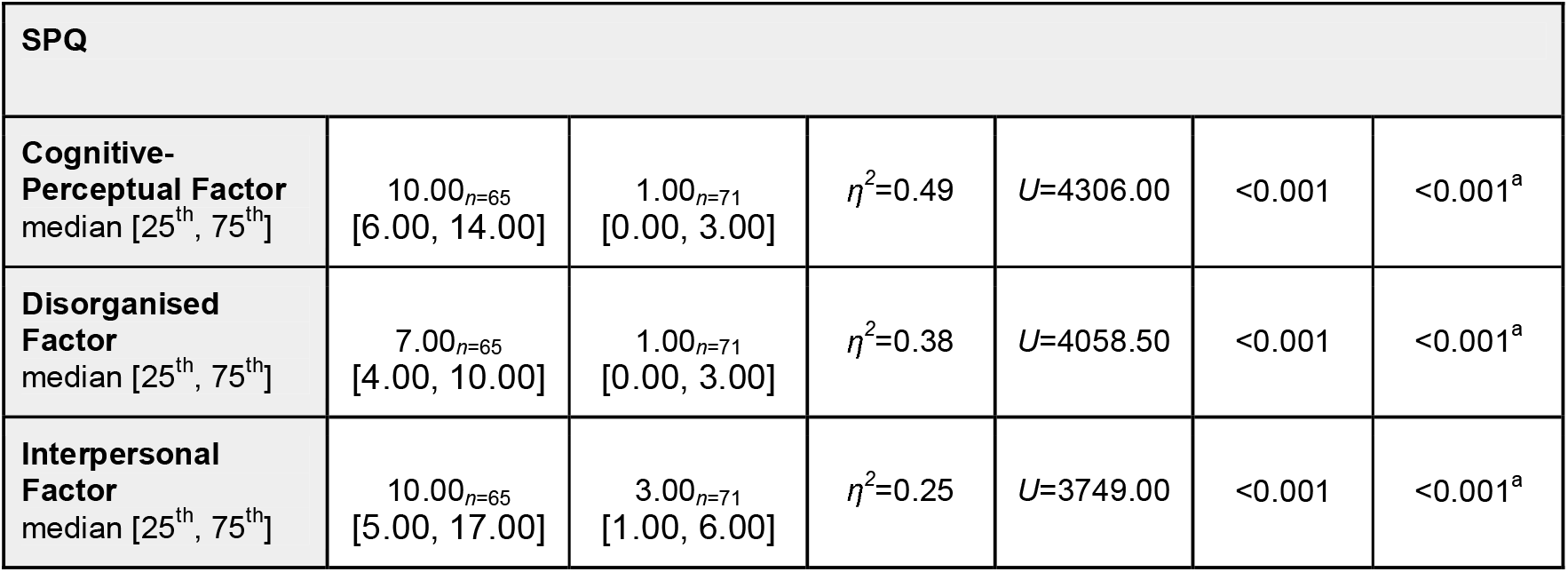
Demographic and preclinical characteristics. **HS** High schizotypy group. **LS** Low schizotypy group. ^a^Benjamini-Hochberg-adjusted *p*-values for *n*=7 comparisons across short Oxford-Liverpool Inventory of Feelings and Experiences (**sO-LIFE**) and Schizotypal Personality Questionnaire (**SPQ**) subscales.

### Assessment of pre-clinical symptoms

Pre-clinical symptoms were assessed with the sO-LIFE questionnaire, including positive schizotypy with the *Unusual Experiences*, disorganised schizotypy with the *Cognitive Disorganisation*, negative schizotypy with the *Introvertive Anhedonia*, and impulsivity with the *Impulsive Nonconformity* subscale, respectively. We assessed convergent validity with the *cognitive-perceptual, disorganised* and *interpersonal* factors (44,45) of the Schizotypal Personality Questionnaire (SPQ) (44,46). SPQ data was missing for *n*=1 HS and *n*=1 LS.

#### Statistical analyses of demographics and pre-clinical symptoms

Demographic (age, sex, handedness, education, BMI, physical activity, alcohol, smoking, cannabis, verbal IQ) and preclinical variables (sO-LIFE subscales and SPQ factors) were compared between HS and LS groups. Continuous variables were analysed using independent-samples *t*-tests, if normality assumptions were valid (assessed with Shapiro–Wilk tests), or analysed with Mann-Whitney U tests otherwise. Categorical variables were compared using chi-square tests. Effect sizes are reported as Cohen’s *d* for *t*-tests, η^*2*^ for Mann-Whitney U tests, and Cramér’s *V* for chi-square analyses. To account for multiple testing across pre-clinical scales, *p*-values were adjusted for *n*=7 (4 sO-LIFE subscales + 3 SPQ factors) comparisons using the Benjamini–Hochberg procedure (47).

### (f)MRI acquisition

Structural and functional MRI images were collected using a 3T MRI system (Magnetom Prisma, Siemens Healthcare, Erlangen, Germany) with a 20-channel radio frequency head coil. Anatomical images were collected with a T1-weighted magnetisation prepared rapid acquisition gradient (MPRAGE) sequence (field-of-view: 256×256×176, resolution: 1mm^3^; TR: 2000ms; TE: 3.37ms; flip angle: 8°; bandwidth: 200Hz/pixel). Functional rs-fMRI was acquired using interleaved T2*-weighted EPI using the following parameters: 36 axial slices, slice thickness: 3.0mm, inter-slice gap: 1.0mm (33% distance factor), field-of-view: 256×256×144mm, voxel size: 3.0×3.0×3.0mm, TR: 3000ms, TE: 28ms, flip angle: 82°, and bandwidth: 2326Hz/pixel. One hundred sixty-eight volumes were acquired (total acquisition time: 8min 24s). Two participants could not be scanned due to safety concerns regarding potential metallic splinters in the eyes that were not disclosed during initial screening and one participant was excluded due to an incidental finding resulting in a final sample of N=138 participants (HS: n=66, LS: n=72). Please, see Supplementary Figure 1 for an overview of the analysis pipeline.

### (f)MRI preprocessing

Preprocessing of the rsfMRI data was conducted using fMRIPrep v23.0.2 (48), which is based on Nipype v1.8.5 (49) following the steps outlined in (50). Briefly, anatomical T1-weighted images underwent intensity non-uniformity correction with N4BiasFieldCorrection (51) of ANTs v2.3.3 (52), followed by skull-stripping and tissue segmentation into gray matter, white matter, and cerebrospinal fluid using FAST (53) of FSL v6.0.5.1:57b01774 (54). Nonlinear spatial normalization to MNI152NLin2009cAsym template space was performed using antsRegistration with brain-extracted versions of both T1-weighted reference and T1-weighted template (MNI152NLin2009cAsym) (55).

For functional data, a reference volume was generated and skull-stripping was performed. Head motion parameters were estimated using *mcflirt* (56), and slice timing correction was applied. Susceptibility distortions were corrected using a fieldmap-less approach (57). Functional images were then co-registered to the anatomical reference using boundary-based registration with *flirt* (58), and normalized to MNI space (MNI152NLin2009cAsym) (55).

### Proton Magnetic Resonance Spectroscopy acquisition & processing

Proton magnetic resonance spectroscopy (^1^H-MRS) acquisition and preprocessing is reported in the Supplement. We consider these analyses to be exploratory due to the voxel placement in the left hippocampus outside of the modelled network.

### Dynamic causal modelling

We estimated directed (effective) connectivity between brain regions and local excitability of different brain regions with spectral dynamic causal modelling (DCM) (59–61). Spectral DCM characterises the low⍰frequency (<0.1□Hz) cross⍰spectral density of spontaneous neuronal activity by estimating the amplitude and the spectral exponent capturing the covariance structure of the hidden neuronal states within each region of a predefined network (61). Because this approach uses a power⍰law form, it provides a deterministic and computationally efficient model that still reflects stochastic neural fluctuations. Connectivity strengths are expressed in Hz, meaning that higher values correspond to faster induced responses in downstream areas. The resulting neuronal dynamics are then transformed into predicted fMRI signals using a haemodynamic forward model (62). All following analyses were performed using the open-source software Statistical Parametric Mapping (SPM12; v12.6; Wellcome Trust Centre for Neuroimaging, London, UK, https://www.fil.ion.ucl.ac.uk/spm) in MATLAB (R2024a for model fitting and R2019b for sensor-level and group-level analyses; The MathWorks Inc., Natick, MA; https://www.mathworks.com).

#### Network selection

In line with Adams et al. (16), we modelled effective connectivity in a 6-region network activated during MMN paradigms (Figure 1), comprising bilateral primary auditory cortex (A1), superior temporal gyrus (STG), and inferior frontal gyrus (IFG) derived from the Glasser parcellation (63) using the glasser_MNI152NLin2009cAsym_labels_p20.nii from (64); see Supplementary Table 1 for Glasser atlas region labels). The reasons for using only these regions are: i) they are likely activated by the (unpredictable) noises happening during an fMRI scan, ii) they involve areas that are implicated in schizophrenia (auditory and frontal areas and connections between them), and iii) we were replicating the methods used previously by Adams et al. (16). We created masks for all six regions using nearest neighbour interpolation to avoid blurring region boundaries. Parcellation masks were resampled to the resolution of the functional imaging data and weighted by grey matter probability in that region to remove spurious white matter and cerebrospinal fluid signals. We used an extraction threshold of 0.5 (i.e. >50% probability of a voxel being grey matter).

**Figure 1.**
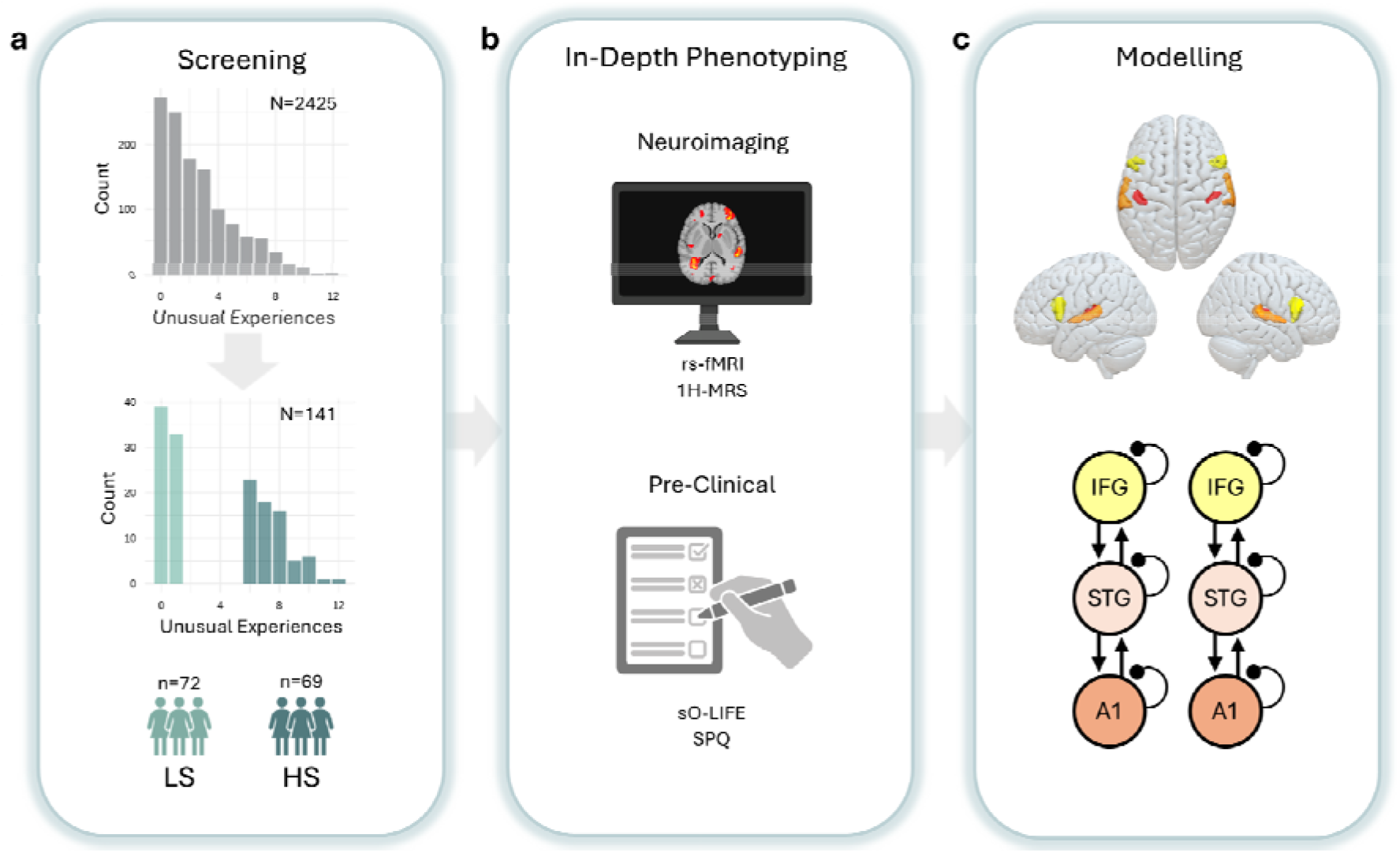
Study Overview. (**a**) Participants from the general population were screened using the short Oxford-Liverpool Inventory of Feelings and Experiences (**sO-LIFE**). Individuals at the extreme ends of the *Unusual Experiences* subscale were assigned to the low (LS; *Unusual Experiences* ≤1) or high positive schizotypy (HS; *Unusual Experiences* ≥6) group and invited back for in-depth phenotyping (**b**) including neuroimaging and pre-clinical assessments with the sO-LIFE and Schizotypal Personality Questionnaire (**SPQ**). (**c**) Effective connectivity was estimated during resting-state functional magnetic resonance imaging (**rs-fMRI**) with dynamic causal modelling including primary auditory cortex (**A1**, red), superior temporal gyrus (**STG**, orange) and inferior frontal gyrus (**IFG**, yellow). ^**1**^**H-MRS** proton magnetic resonance spectroscopy.

#### First-level GLM

To mirror the analysis of Adams et al. (16), we performed a global signal regression (GSR) using the confound regressors recommended by *fMRIprep* (https://fmriprep.org/en/latest/outputs.html#confounds). We used a total of 36 regressors, following previous guidelines (65,66). These include six motion regressors, which are estimated during the *fMRIprep* pipeline, three confound regressors (white matter, CSF, global signal), as well as their first-order (temporal) derivatives and quadratic terms accounting for non-linear effects. To filter and remove low-frequency noise components from the data, three discrete cosine basis functions were included in the first-level general linear model (GLM) to remove low-frequency drifts. Additionally, stick regressors computed with thresholds of 0.5mm for frame displacement (FD) and 1.5 for standardised DVARS were used to model outlier trials as done previously (50). For each participant, we extracted time-series data from six predefined regions of interest.

To test for differences in model fit across groups, R^2^-values were first fisher-r-to-Z transformed, but since normality assumptions were still not met, we proceed with Mann-Whitney U tests. The model fit was R^2^=59.00% [51.29%, 65.56%] (median [25th, 75th]) and did not differ significantly across groups (HS: R^2^=58.54% [51.38%, 66.92%], LS: R^2^=59.68% [51.07%, 64.18%]; η^*2*^*<0*.*01,U*=2486.00, *p*=0.641; Figure S2a).

#### Second-level GLM

To take not only connectivity parameter estimates but also their uncertainty into account, we tested for second-level (between-subject) effects on synaptic parameters with a Bayesian GLM (67,68). In this parametric empirical Bayes (PEB) approach, the maximum a posteriori (MAP) connectivity parameter estimates from the first-level DCMs are taken to the second (group) level where they are used as the dependent variable in a Bayesian GLM to be explained by variables of interests (group or pre-clinical symptoms) and covariates. Correlation analyses with pre-clinical symptoms were restricted to the HS group as the LS group showed overall low pre-clinical scores and limited variance.

Using PEB, we compared models with different combinations of forward, backward and local excitability modulations (Figure S2b) as done previously (16). Since there was no clear winning model (Figure S2c), we performed Bayesian model averaging over the resulting parameter estimates. Since groups were prospectively matched on key covariates and did not significantly differ on any of the other recorded covariates (Table 1), no covariates were included in the second-level GLM. We report Bayesian posterior probabilities (P), which quantify the probability that a given effect is present (i.e., non-zero), given the data and model. Analogous to the frequentist α=0.05, we interpret P>.95 as strong evidence for an effect (69) and report beta estimates in logspace and their 95% Bayesian confidence intervals as a measure of effect size.

For example, a second-level PEB group effect of 0.15 corresponds to e^0.15^≈1.162, i.e. about 16% increased connectivity.

## Results

### Demographic and pre-clinical characteristics

Participants did not differ in terms of demographic characteristics (Table 1). HS scored significantly higher on all sO-LIFE and SPQ subscales (all *p*<0.05).

### Effective connectivity in individuals with high vs low schizotypy

HS showed significantly reduced backward connectivity from IFG to STG bilaterally (both P>0.95; Figure 2a) and increased backward connectivity from left STG to A1 (P>0.95; Figure 2a).

**Figure 2.**
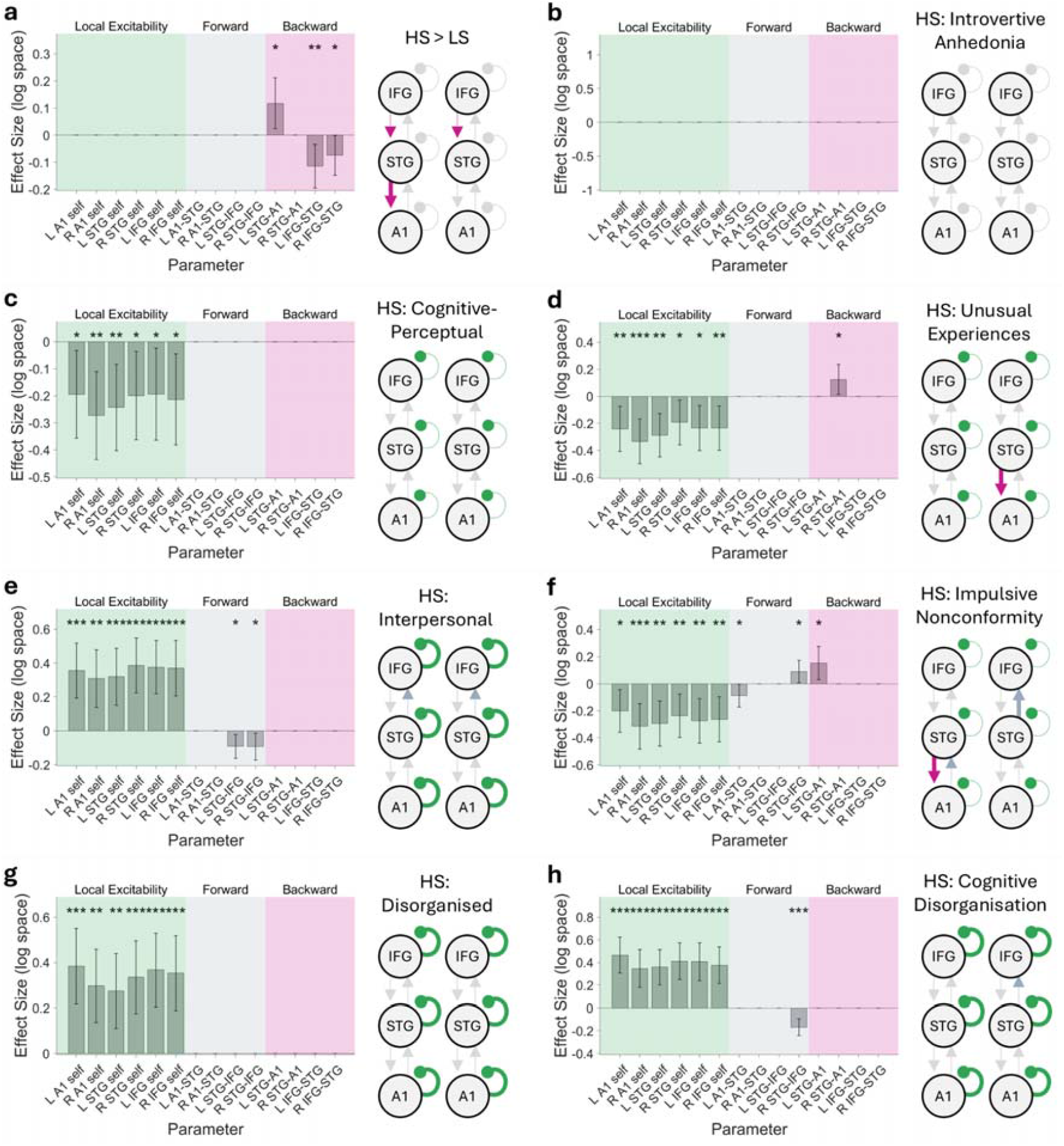
Group differences in effective connectivity correlations between effective connectivity and pre-clinical symptoms. (**a**) Changes in effective connectivity (in log space) in individuals with high (**HS**) compared to low positive schizotypy (**LS**). In the model diagram, reduced connectivity is illustrated with dotted lines. Across all panels, bar height represents the mean (maximum a posteriori) and error bars indicate the 95% Bayesian confidence intervals. Correlations between effectivity connectivity (in log space) and the (**b**) introvertive anhedonia (**d**) unusual experiences, (**f**) impulsive nonconformity, or (**g**) cognitive disorganisation subscales of the Oxford-Liverpool Inventory of Feelings and Experiences (**sO-LIFE**) and the (**c**) cognitive-perceptual, (**e**) interpersonal, (**g**) disorganised factor of Schizotypal Personality Questionnaire (**SPQ**) in the HS group. In model diagrams, negative and positive correlations are illustrated with dotted or thick lines, respectively. Note that local excitability (green) is modelled through inhibitory self-connections; thus reduced (**c, d, f**) and increased self-connectivity (**e, g, h**) model either hyper- and or hypoexcitability of a given region, respectively. Asterisks indicate a significant group effect (a) or correlation (b-f) with a posterior probability of (*) P>.95, (**) P>.99 or (***) P>.999.

In exploratory MRS analyses in the HS group (see Supplement), we found a positive correlation between left IFG-STG backward connection and hippocampal E/I balance as approximated through Glutamate(Glu)/Gamma Aminobutyric Acid (GABA) ratio measured with ^1^H-MRS (P>0.999, Figure S3a). This effect was driven by both negative correlations with GABA (P>0.999, Figure S3b) and positive correlations with Glu (P>0.95, Figure S3c).

### Correlations with pre-clinical symptoms

#### Unusual Experiences

Within the HS group, increased right STG-to-A1 backward connectivity and reduced self-connectivity across all regions (i.e., increased excitability or disinhibition) was correlated with higher *Unusual Experiences* scores (all P>0.95; Figure 2d). We found convergent findings with the SPQ suggesting that reduced self-connectivity (increased excitability) across all regions was correlated with higher scores on the *cognitive-perceptual* factor (all P>0.95; Figure 2c).

#### Introvertive Anhedonia

Within the HS group, there was no significant correlation between effective connectivity and *introvertive anhedonia* (Figure 2b).

#### Cognitive Disorganisation

Within the HS group, reduced right STG-to-IFG forward connectivity and increased self-connectivity across all regions (i.e., reduced excitability) correlated with higher *Cognitive Disorganisation* scores (all P>0.95; Figure 2h). We found convergent validity with the SPQ, suggesting increased self-connectivity across all regions (i.e., reduced excitability) was associated with higher *disorganised* factor scores (all P>0.95; Figure 2g).

#### Impulsive nonconformity

Within the HS group, reduced self-connectivity across all regions (i.e., increased excitability) correlated with increased *Impulsive Nonconformity* scores (all: P>0.95; Figure 2f). Higher SPQ *interpersonal* factor scores correlated with the connections (i.e., reduced self-connectivity or increased excitability across all regions) and with bilaterally reduced STG-to-IFG forward connectivity (all: P>0.95; Figure 2e). Additionally, reduced left A1-to-STG and increased right STG-to-IFG forward connectivity as well as increased left STG-to-A1 backward connectivity correlated with higher *Impulsive Nonconformity* (all: P>0.95; Figure 2f).

## Discussion

This study investigated changes in effective connectivity and cortical excitability across individuals with high compared to low positive schizotypy. There are several key results of this study. First, we found that individuals with high positive schizotypy had reduced IFG-to-STG backward connectivity compared to those with low schizotypy. Second, we found that disinhibition across the auditory cortex-IFG network was associated with more psychotic-like experiences and increased impulsive nonconformity in individuals with high positive schizotypy. Third, we found that reduced excitability in the same network was correlated with stronger cognitive disorganisation.

### Reduced fronto-temporal drive is found across the psychosis spectrum and may reflect reduced E/I balance

We found reduced bilateral IFG-to-STG backward connectivity in individuals with high positive schizotypy. Adams et al. (16) found reduced excitability in IFG when modelling the same network of regions during rs-fMRI in patients with chronic schizophrenia. Our results are similar to their findings in that we found reduced backward connectivity in our study vs reduced local excitability in theirs—resulting in reduced top-down information flow from frontal cortex to temporal regions. Our results thus support the view that frontal cortex dysfunction may be a key biological mechanism cutting across the psychosis spectrum extending to individuals from the general population with high positive schizotypy.

Interestingly, reductions in the right fronto-temporal backward connection were also observed in individuals with 22q11.2 deletion syndrome during a MMN paradigm (70). Bilateral reductions in backward connectivity were also found to correlate with psychotic-like experiences assessed with the community assessment of psychic experience (71) in a pooled sample of 22 healthy controls, 20 schizophrenia spectrum patients and 20 nonpsychotic inpatients (72). However, others have found increased and not reduced fronto-temporal backward connectivity in early illness schizophrenia patients in the left hemisphere (73) or the right hemisphere of chronic patients with schizophrenia spectrum disorders (74) (see Gütlin et al (75), for a recent narrative review). These incongruent results are perhaps due to the small sample sizes in some of these studies (e.g., Dima et al. (73) only included *n*=12 patients). To the best of our knowledge our study is the largest single-centre study of individuals with high schizotypy to date.

More recent larger studies including ours appear to converge on finding reduced E/I ratio across the psychosis continuum (16,20,29–31). In line with this, we found reduced left IFG-to-STG backward connectivity to be correlated with reduced E/I balance in hippocampus, which was driven by changes in both Glu and GABA levels as measured with ^1^H-MRS (Figure S3). Although the voxel placement in the left hippocampus precludes us from drawing any strong conclusions, if hippocampal E/I balance correlates with E/I balance in other cortical areas, this finding may suggest that reduced fronto-temporal drive could reflect reduced excitatory drive and thus reduced E/I balance. Future studies perhaps using whole brain MRS – are needed to clarify the relationship between hippocampal and prefrontal glutamate levels.

### Allostatic cortical disinhibition may drive psychotic-like experiences

Interestingly, we found that widespread disinhibition was associated with more severe psychotic-like experiences, while the top-down fronto-temporal drive was reduced overall. This seemingly paradoxical result has been found across neuroimaging modalities (EEG, fMRI) (16) and clinical populations (CHR-P, chronic patients) (16,30). A recent mega-analysis of MRS studies also found reduced glutamate levels in patients with schizophrenia but a positive correlation between glutamate and positive symptom severity (35). This result suggests that allostatic rebalancing of a primary glutamatergic deficit – as recently theorised (10) and supported by post-mortem data (76) – may drive psychotic symptoms. This relationship has been seen in CHR-P (30) and chronic patients (16). Importantly, our results corroborate this interpretation and suggest that the same biological mechanism may underlie positive schizotypy in the general population. We found that widespread cortical disinhibition was also associated with increased impulsivity, similar to positive symptom severity. This suggests that cortical disinhibition may be a shared biological mechanism of positive symptoms and impulsivity.

### Reduced cortical excitability relates to disorganised symptoms

Interestingly, we found reduced excitability correlated with greater cognitive disorganisation. This finding is consistent with a large study investigating excitation/inhibition imbalance across transdiagnostic psychosis biotypes, which found that reduced pyramidal cell excitability correlated with more severe cognitive symptoms in patients (29,31). One speculative explanation is that reduced excitability results in shallower attractor states, which have been postulated to underlie reasoning biases (77,78) and working memory deficits in patients with schizophrenia (79).

### Clinical implications

Fronto-temporal backward connectivity may be used as a biomarker to enrich psychosis risk prediction models along with demographic (80), structural magnetic resonance imaging (81–83) or EEG measures (84,85) or as part of a sequential test battery as suggested previously (86–88). Cortical excitability may be a treatment target to reduce positive symptoms and impulsivity.

### Limitations and future directions

A few limitations of this study merit attention. First, the MRS voxel was positioned over the left hippocampus, which is not part of the MMN network. We therefore consider these results to be exploratory. Although hippocampus measures may be a proxy for overall E/I imbalance in the brain, future research should investigate the MMN network using high-field MRI. Second, the reliability of effective connectivity parameters needs to be assessed, especially if parameter estimates are used as biomarkers for prognosis or target engagement (89,90). Third, future studies should examine the pharmacological basis of fronto-temporal backward connectivity using pharmacological studies and test whether interventions including brain stimulation targeting cortical excitability reduce impulsivity or positive symptoms as suggested by recent findings (91).

## Conclusion

This study found evidence that reduced frontal cortex function represents a biological mechanism that cuts across the psychosis continuum to preclinical populations. Disinhibition across the auditory cortex-IFG network was associated with more psychotic-like experiences and increased impulsive nonconformity and reduced excitability in the same network was correlated with stronger cognitive disorganisation.

## Code availability

The analysis code will be made publicly available at https://github.com/daniel-hauke/dcm_grace upon acceptance of the paper.

## Data availability

Access to the data can be arranged through a formal collaboration with AS. Contact AS to inquire about such collaborations.

