## Supplement for "Fronto-Temporal Dysconnectivity and Cortical Excitability in High Schizotypy: Associations with Symptom Dimensions"

† Drs. Adams and Schmidt are joint last authors.

### Supplementary Methods

#### Proton Magnetic Resonance Spectroscopy acquisition & processing

During the same session as the structural and functional MRI data acquisition we conducted a proton magnetic resonance spectroscopy (1H-MRS) where spectra of the left hippocampus were obtained using water-suppressed Mescher-Garwood point resolved spectroscopy (MEGA-PRESS). Due to unavailable unsuppressed water MEGA-PRESS spectra, Gamma Aminobutyric Acid (GABA) and Glutamate (Glu) were scaled to *N*-acetylaspartate (NAA) for signal referencing. Spectra with a Cramér-Rao lower-bound >20% were excluded from analysis. For more details, see Iseli et al. (2025).

| Region | Region Name (Glasser) | Region Number (Glasser) |
|---|---|---|
| Left primary auditory cortex (lA1) | L_A1_ROI | 24 |
| Right primary auditory cortex (rA1) | R_A1_ROI | 204 |
| Left superior temporal gyrus (lSTG) | L_A4_ROI | 175 |
| Right superior temporal gyrus (rSTG) | R_A4_ROI | 355 |
| Left inferior frontal gyrus (lIFG) | L_44_ROI | 74 |
| Right inferior frontal gyrus (rIFG) | R_44_ROI | 254 |

**Supplementary Table 1**. **Region labels and numbers from the Glasser parcellation**.

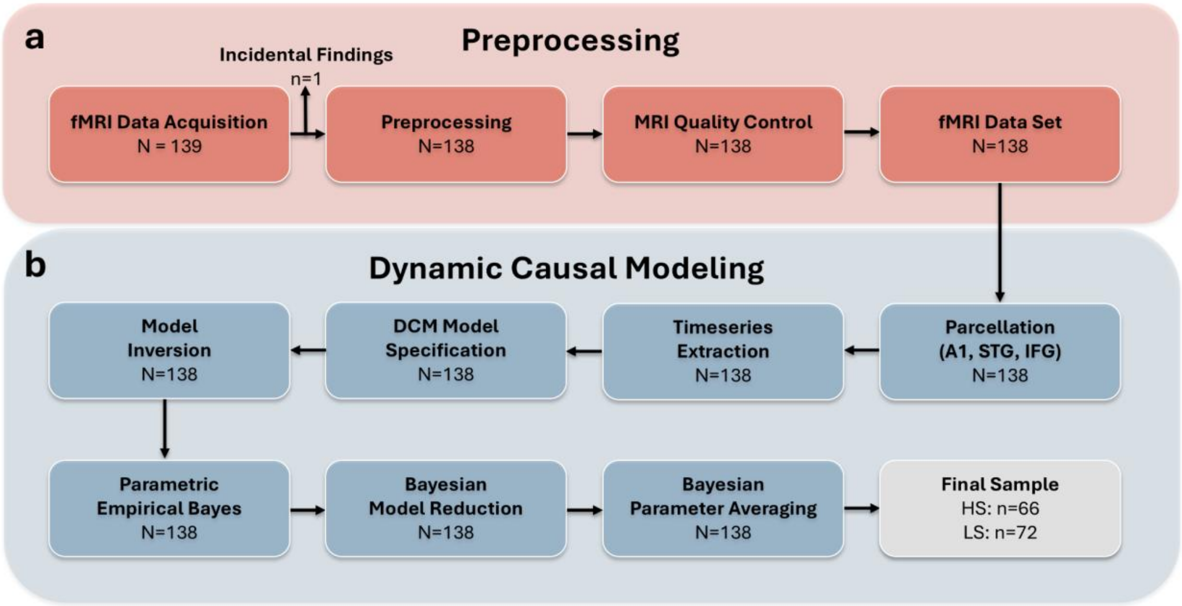


**Figure S1. Data processing and analysis pipeline.** Schematic overview of the three-stage analytical workflow with preprocessing displayed in (**a**), time series extraction in (**b**) and dynamic causal modelling pipeline in (**c**). **LS** Individuals with low positive schizotypy. **HS** Individuals with high positive schizotypy.

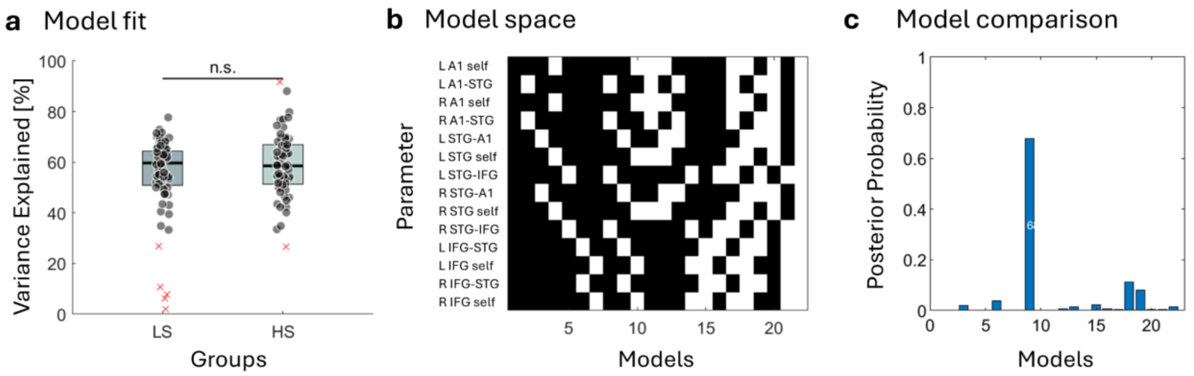


**Figure S2. Model fit, model space and Bayesian model comparison results.** (**a**) Model fit across groups. Model fit measured as variance explained ($R^2$) across groups is shown. Box boundaries represent the 25th and 75th quantiles, with the solid horizontal line indicating the median. Whiskers extend to 1.5 times the interquartile range. (**b**) Model space. Each row represents a model containing different combinations of extrinsic or self-inhibitory connection parameters (in white). (**b**) Model comparison results. The posterior probability of each model is shown, following Bayesian model reduction. There was no clear winning model. Bayesian parameter averaging across all models was performed to investigate the parameter effects reported in the main manuscript.

### Supplementary Results

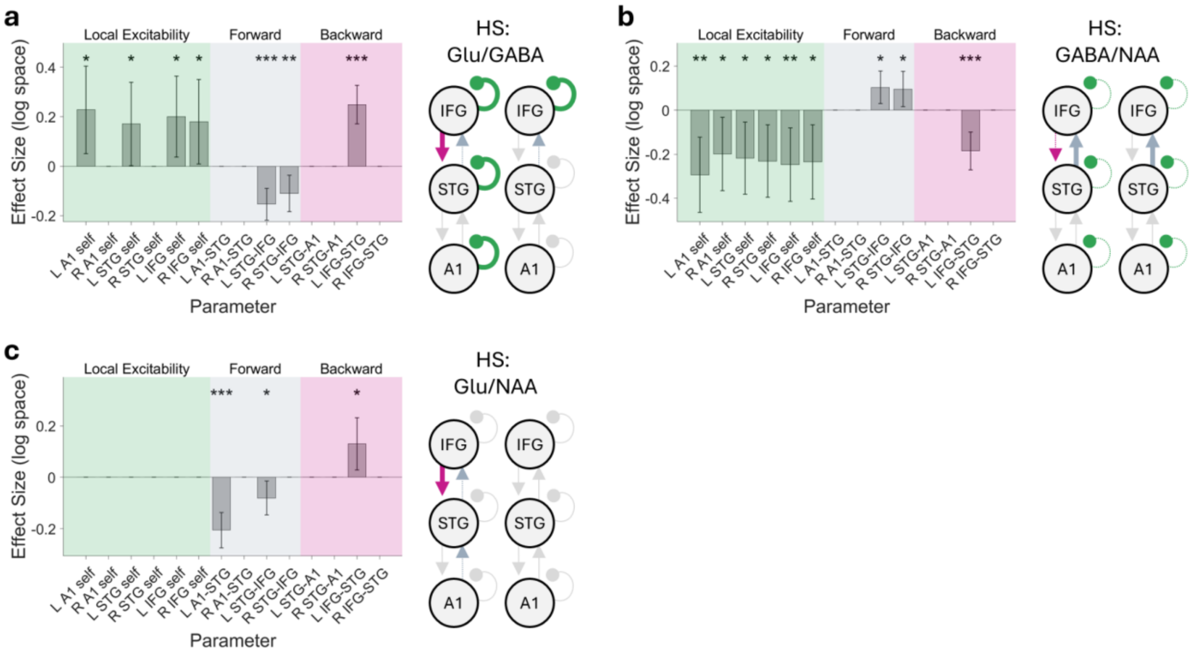


**Figure S3. Correlations between effective connectivity and hippocampal excitation/inhibition balance.** Correlations between effectivity connectivity (in log space) and (**a**) hippocampal excitation/inhibition balance measured as the GABA/Gl ratio (**b**) hippocampal GABA and (**c**) hippocampal glutamate levels. in individuals with high schizotypy (**HS**). In model diagrams, negative and positive correlations are illustrated with dotted or thick lines, respectively. Note that local excitability (green) is modelled through inhibitory self-connections; thus reduced (**b**) and increased self-connectivity (**a**) model either hyper- and or hypoexcitability of a given region, respectively. Asterisks indicate a significant correlation with a posterior probability of (*) P>.95, (**) P>.99 or (***) P>.999. **GABA** Gamma Aminobutyric Acid. **Glu** Glutamate. **NAA** N-acetylaspartate.
